# Geometric probability, psychophysics and invariance

**DOI:** 10.1101/531889

**Authors:** Yuval Hart, L. Mahadevan

## Abstract

The perception of the noisy visual world around us naturally combines geometry and probability with psychophysics. So how do we perceive geometric objects from a probabilistic perspective, i.e. infer randomness in a spatial setting ? To test this psychophysically, we use a set of simple experiments to distinguish between probability distributions of planar line images connected with Buffon’s needle and Bertrand’s paradox, two classic exemplars of geometric probability. We find that participants associate greater randomness with images that are invariant under the sub-groups of translation, rotation, and scale. An information theoretic framework centered around the Radon (Hough) transform captures the observed behavioral results, and suggests that symmetry and chance are embedded in human visual perception.

---

The perception of the visual world has at its core both geometry and probability; geometric inference allows us to assess distances and directions in the world, while probabilistic inference allows us to assess the odds of an outcome. While the former are universal[1–3], the latter are often biased by priors and heuristics that arise due to limited resources [4–9].

In visual perception, previous studies focused on mechanisms for local texture inference. Among which are: first-order, second-order, and higher-order intensity statistics [10–12], the well-known filter-rectifier-filter models [13–15], and more recently, texture flows mechanism [16, 17] which preserves local curvatures. Recent work complements this by showing that people can distinguish between ordered and disordered images with discrete symmetries [18, 19], and a statistical physics perspective helps to explain the ability to discriminate among images. Here we consider if and how humans can distinguish patterns that exhibit the breaking of continuous symmetries associated with simple planar Lie groups in noisy settings, by considering the conjunction of randomness, probability, and visual perception.

To explore this avenue quantitatively, we use psychophysical experiments inspired by the framework of geometric probability, the generalization of probability problems to spatial settings [20]. It relies on defining natural probabilistic measures that are invariant to transformation groups such as translation, rotation, and scale [21–23]. Two of the oldest problems in geometric probability are the following: Buffon’s needle, which asks the following question [24]: what is the chance that a randomly oriented line of given length *L* intersects a fixed parallel grid of lines with a given spacing *l*(*> L*) (Fig 1a). The answer, 2*L/πl*, follows from simple geometric considerations assuming that the orientation distribution is uniform. Bertrand’s paradox [25] asks what is the probability that a random chord on a circle is longer than a side of an equilateral triangle inscribed in the circle? Bertrand showed how different random processes yield different answers to the same question, and used this example to argue that the notion of randomness is ill-posed in the absence of a process that generates the randomness, i.e., in the absence of a well defined probabilistic measure. Later, Poincare [26] pointed out the link between different realizations of randomness and the symmetries they hold and Jaynes [27] built on this by arguing that the assessment of randomness should be dictated by the principle of ‘maximum ignorance’, i.e. the method which holds as many invariances as possible (rotation, translation and scale) is to be preferred.

**FIG. 1.**
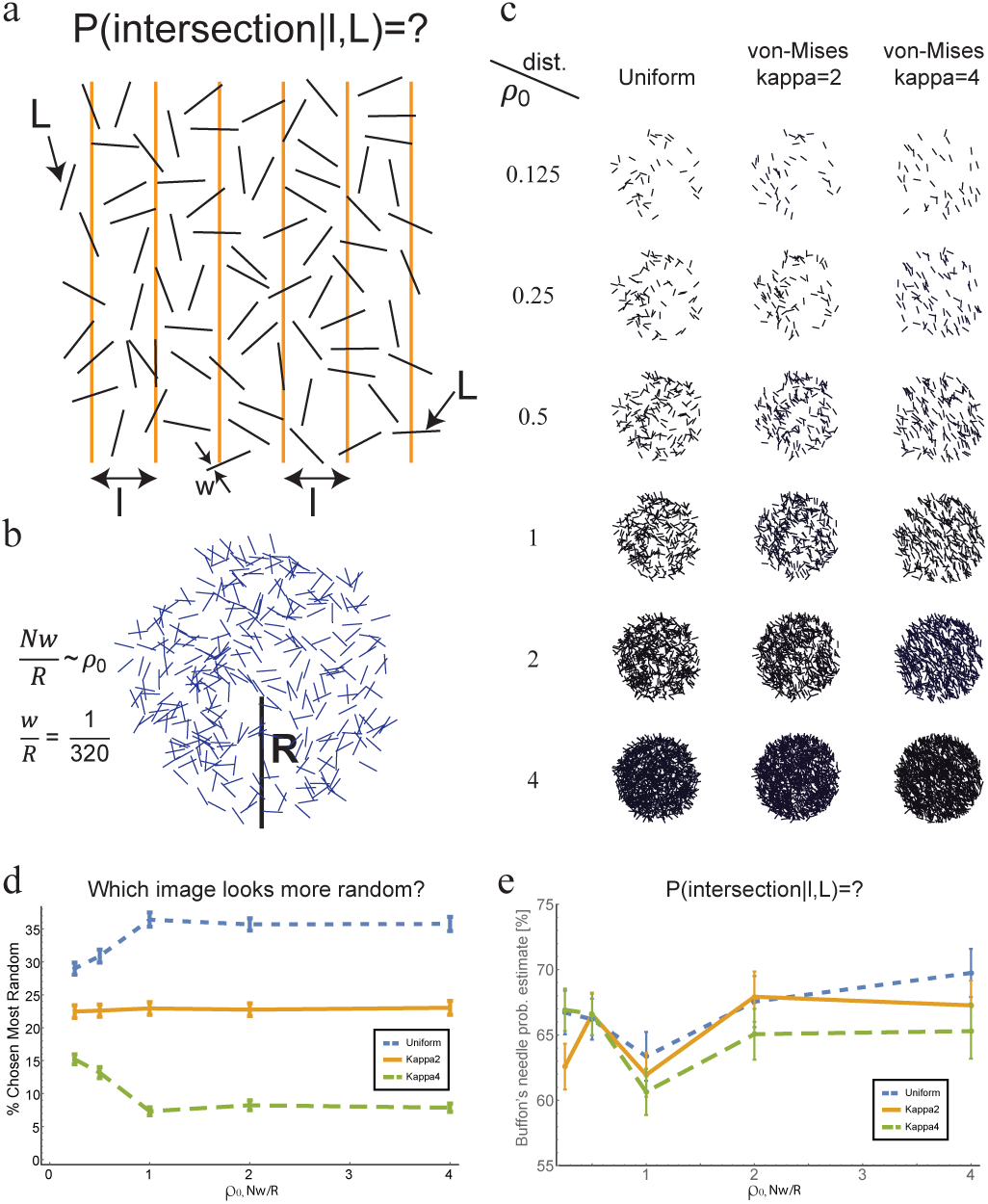
Psychophysics of Buffon’s needle. **a)** Buffon’s needle problem: Given a needle of length *L* dropped on a plane ruled with parallel lines *l* units apart, what is the probability that the needle will cross a line? **b)** In the psychophysical version of the problem, images shown consist of a number *N* of random lines inside a circle of radius *R*, with varying density is *ρ*_0_ = *Nw/R*, where *w* is the line width. **c)** Example of images with different *ρ*_0_ values from a von-Mises distribution 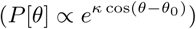 with *κ* = 0, 2, 4. **d)** Participants’ choices for the most random images as a function of *ρ*_0_. We see that participants choose *κ* = 0 (blue, short dash) as the most random, and then *κ* = 2 (green, long dash), and *κ* = 4 (gold, solid) as the least random. **e)** Participants’ estimates are close to the correct value 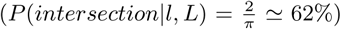. Distribution 1 (*κ* = 0) in blue, Distribution 2 (*κ* = 2) in gold, and Distribution 3 (*κ* = 4) in green.

To convert Buffon’s needle problem to a visual psychophysical task, we use a variant of the problem that presents the simultaneous realizations of multiple lines. Our stimuli take the form of lines of identical length drawn from a given orientational distribution (depicted within a circular domain to withhold further orientational cues from the image boundary). In a perceptual setting we need to move beyond the abstraction of infinitesimal points and lines to account for their finite size and width. In the case of lines in a circle, a natural parameter is the density of coverage, *ρ*_0_ = *Nw/R*, where *N* is the number of chords, *w* is the width of each chord, and *R* is the radius of the circle. When *ρ*_0_ *→* 0 or *ρ*_0_*→ ∞*, the image corresponds to a full disc of either all white or all black, so that there is no information in the image; however, for intermediate values of *ρ*_0_ (Fig 1b,c) there is directional information in the image. To quantify this, we use the von-Mises distribution [28], *P* (*θ*) = exp[*κ* cos(*θ-θ*_0_)]*/I*_0_(*κ*), the analog of the Gaussian distribution of a periodic random variable with the mean angle (*θ*_0_) and the variance (1*/κ*). We used three values of *κ* associated with *κ* = 0 for Distribution 1, *κ* = 2 for Distribution 2, and *κ* = 4 for Distribution 3 (Fig 1b,c).

We asked participants (N=470, online experiment, see Methods) to identify the odd-one-out among four images, three of which were taken from the same orientation-sampling distribution and one from another distribution (see Methods). Participants differentiated well between Distribution 1 (uniform) and Distribution 3 (von-Mises, *κ* = 4), with accuracy increasing with density *ρ*_0_ (see SI, Fig S1). Participants distinguished only slightly above chance level (but significantly for most of the density range) between Distribution 2 (*κ* = 2) and the two other distributions (uniform, von-Mises with *κ* = 4, see SI and Fig S1). Next, we asked participants to choose the more random image among two images created using two different distributions chosen from the above three. Interestingly, randomness estimates correlated with the level of rotational symmetry in the image: participants chose Distribution 1 (uniform) as most random, followed by Distribution 2 (*κ* = 2) and lastly Distribution 3 (*κ* = 4). Furthermore, the ordering of the distributions increased with density up to *ρ*_0_ = 1 and then saturated (Fig 1d). To validate the participants quantitative assessment of randomness, in a separate task we asked participants to review single images from each distribution, and score their randomness between 0 (ordered) to 100 (maximally random) with similar results (see Methods, and Fig S2). Finally, we asked participants to estimate the probability that a random line intersects a grid of lines with equal spacing between them (Buffon’s needle probability estimate). We find that the mean of participants collective estimates resulted in probability estimates which were close to the correct probability [20, 24] (2*/π*, see Fig. 1e). All together, these results show that participants are able to discern the connection between symmetry (in this specific case, orientational order) and chance (randomness) in a simple setting associated with a realization of Buffon’s needle.

While Buffon’s needle explores the consequences of rotational invariance, it is agnostic to translational and scale symmetry. To explore if and how we are able to estimate randomness and probability associated with translational and scale invariances in planar images of lines, we turn to Bertrand’s paradox [20, 25] that begins with the question: what is the chance that a random chord of length *L* is longer than a side of an equilateral triangle of length *l* inscribed in a circle, i.e. *P* (*L > l*) (see Fig 2a)? Bertrand suggested three different answers, relying on different processes associated with the choice of a random chord in the first method, denoted *C*, a chord is chosen by connecting two points chosen from a uniform distribution on the circumference of the circle; the resulting probability is *P*_*C*_ (*L > l*) = 1*/*3 (see Fig 2a). This method is rotational invariant, but not scale or translational invariant. For the second method, denoted *R*, one chooses a radius drawn from a uniform distribution of orientations, and then the midpoint of the chord is drawn from a uniform distribution along the radius; the resulting probability is *P*_*R*_(*L > l*) = 1*/*2. This method is both rotational, translational, and scale invariant. For the third method, denoted *A*, one chooses the midpoint of the chord drawn from a uniform distribution on the area of the circle (see Fig 2a); the resulting probability is *P*_*A*_(*L > l*) = 1*/*4. This method is both rotational and scale invariant, but not translation invariant. Since only the second method, *R*, with probability *P*_*R*_ = 1*/*2 satisfies rotational, scale and translational invariance [27], the paradox is resolved by linking probability, randomness, and symmetry. But is this true cognitively as well?

**FIG. 2.**
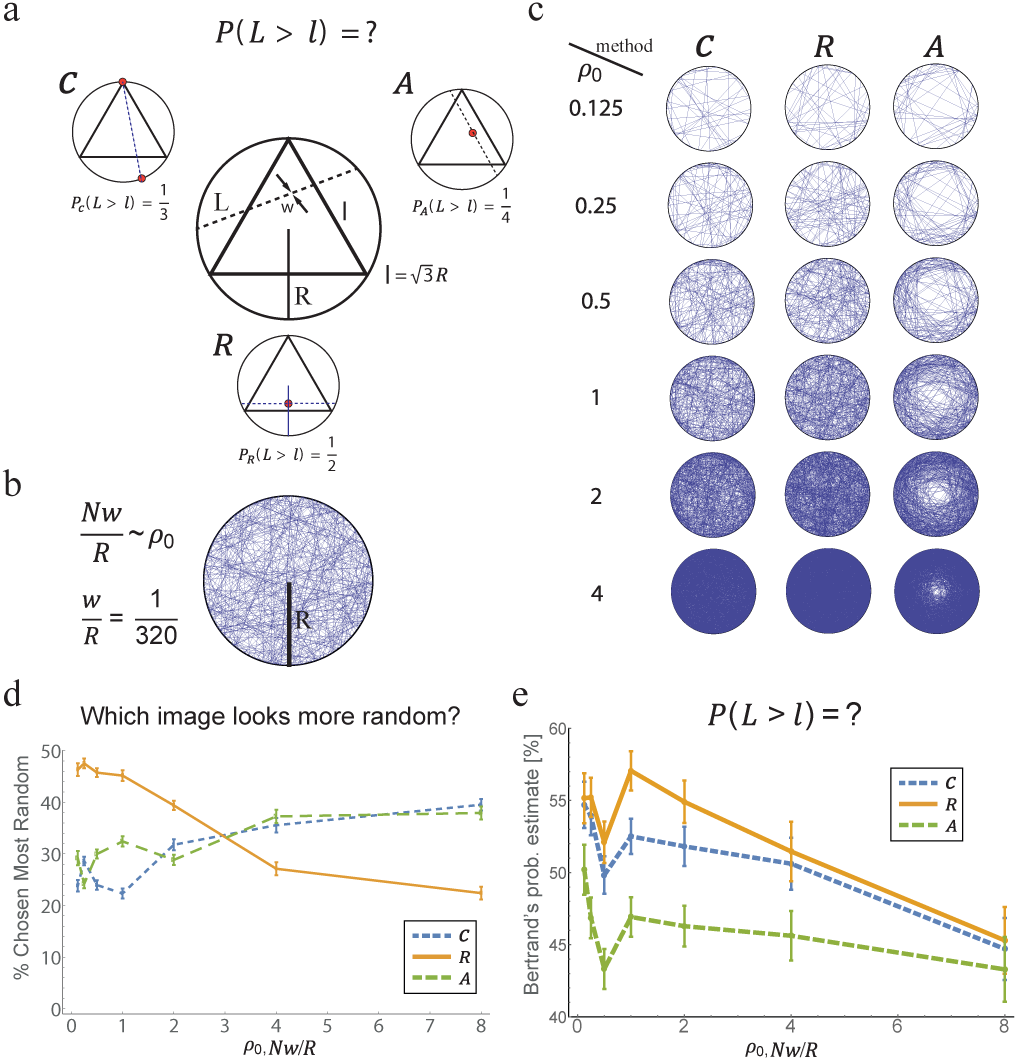
Psychophysics of Bertrand’s paradox. **a)** Bertrand’s paradox: What is the chance that a random chord of length *L* is longer than a side of an equilateral triangle of length *l* inscribed in a circle, i.e. *P* (*L > l*)? Three different methods to draw random chords give different answers: *C* choose two random points along the circumference and draw a chord between them, *P*_*C*_ (*L > l*) = 1*/*3. *R* choose a random radius, choose a random point on the radius and draw a chord with that point as its mid-point, *P*_*R*_(*L > l*) = 1*/*2. *A* choose a random point from the circle’s interior area, that is the random chord’s midpoint, *P*_*A*_(*L > l*) = 1*/*4. **b)** Each image consists of random chords taken from the random processes *C, R, A* with a fixed width of chords and radius. Chords density is given by *ρ*_0_ = *Nw/R*, where N is the number of chords, w the chords’ width and R the radius. **c)** Example of images shown to participants at different *ρ*_0_ values and different methods. **d)** Participants’ choices for the most random method as a function of *ρ*_0_. Each curve represents the fraction of times participants chose a specific method. **e)** Participants estimates of the Bertrand’s problem probability follows the ranking of the correct probabilities in the range of geometrical informative chord densities (0.125 *< ρ*_0_ *<* 3). Method *C* in short-dashed blue line, method *R* in gold, and method *A* in long-dashed green line.

To explore this question, we show participants images of many chords on a circle drawn from the three distributions above (Fig 2b,c). We first asked participants (N=866, online experiment, see Methods) to identify the odd-one-out among four images of different chord distributions, three images taken from the same method and one from a different method. Participants identify the odd-one-out with high accuracy which increases sharply as *ρ*_0_ increases from 0.125 to 1 and saturates (see SI, Figs S3-S4). Next, we asked participants to choose which of two images of chord distributions derived from different methods is more random (see Methods). At the lowest density, *ρ*_0_ = 0.125, method *R*, which satisfies all invariances (rotational, translational and scale) is ranked the most random, whereas methods *C* and *A* (which do not hold all invariances) lag behind (Fig 2d). At *ρ*_0_ ≃2, the perceived randomness associated with method *R* declines and the perceived randomness associated with methods *C* and *A* increases. This suggests an optimal density for assessing randomness (Fig 2d). Similar results were obtained when each method was shown separately and participants were asked to score the randomness of each image (0-not random, 100-fully random). Method *R* scored highest at low chord densities, with a similar transition at *ρ*_0_ ≃2 (see Fig S5). Our results suggest that for most chord densities, participants’ assessment of randomness correlates with the geometrical invariances of chord distributions. Lastly, we asked participants to estimate the probability for each method. We find that participants’ estimates of probability were not accurate in their magnitude (Fig 2e). However, participants estimates of the probability preserved the ranking of the real probabilities, i.e. they assigned *P*_*R*_ *> P*_*C*_ *> P*_*A*_ (see Fig 2e). These differences were significant throughout all chord densities between method *A* and methods *C* and *R* (Mann-Whitney test, *p <* 0.001), and between method *C* and method *R* in the geometrically informative regime (0.5 *≤ ρ*_0_ *≤* 2, Mann-Whitney test, *p <* 0.001).

The ability to estimate randomness and probability associated with invariances raises the question of what perceptual computational mechanism might be at work. The Radon (Hough) transform of the image *f* [29, 30], is a natural measure that is invariant to translation, rotation and scale changes [26]. Defined as 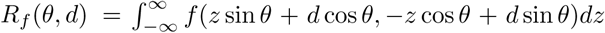 it measures the orientational information content along the line at an angle *θ* and distance *d* from the origin. In Fig. 3a, we show the Radon transform of images associated with the methods *C, R, A* as a function of chord density; the Kullback-Leibler divergence shows that methods *C* and *A* are similar, but methods *R* and *A* are quite different, matching participants’ behavioral results (see SI, Fig S1, and Figs S7-S10).

**FIG. 3.**
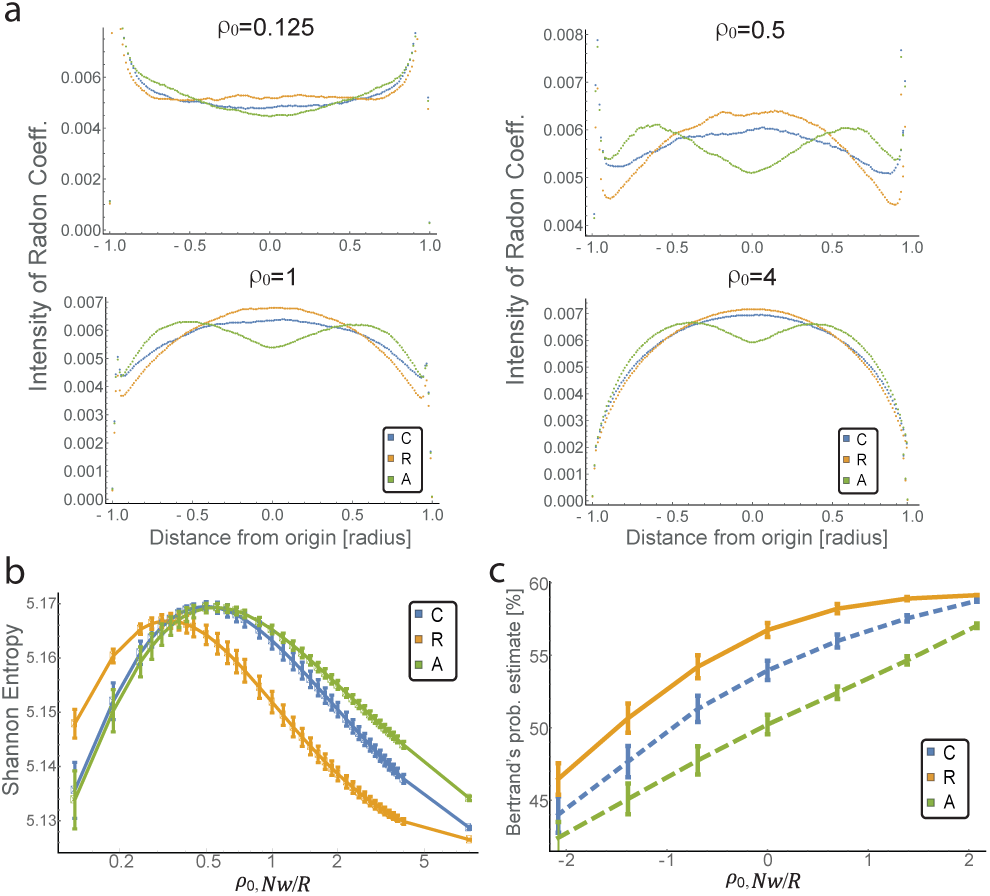
Radon transform captures participants’ behavioral results. **a)** Examples of Radon transform of images with chord densities of: 0.125, 0.5, 1 and 4. x-axis represents distance from origin (as fraction of the radius) and y-axis is the normalized Radon coefficient intensity. *C, R* and *A* methods are represented by blue, gold and green lines respectively. Shannon entropy of the Radon transform curves for all methods. *C, R* and *A* methods are represented by blue, gold and green lines respectively. **c)** The probability estimates of Bertrand’s paradox calculated from the Radon transform of the different methods and density values. The Radon transforms maintains the ranking of the different methods’ probabilities, with similar values to the behavioral estimates (compare with Fig. 2e). Method *R* (gold solid line), method *C* (blue solid line), and method *A* (green solid line). **b,c)** Error bars are standard deviations values.

To capture estimates of randomness we use the correspondence between randomness and Shannon entropy the more bits needed to describe the distribution, the more random it is. In Bertrand’s paradox, the physicality of the problem (density of chords) dictates the amount of information available. Indeed, the Shannon entropy associated with the Radon transform of the images of methods *C, R, A* (Fig 3b) shows non-monotonic behavior, since in the two limits *ρ*_0_*→*0 and *ρ*_0_*→ ∞*, geometrical information is impoverished. Interestingly, similar to assessment of randomness by participants in our experiments, the Shannon entropy of method *R* is larger than that of the two other methods at low chord densities while at high chord densities, the two methods (*C* and *A*) show entropy values higher than method *R*. Moreover, the Shannon entropy curves of methods *A* and *C* coincide, as is the case for participants assessments of randomness in the psychophysical experiments. Similarly, applying the Radon transform on the Buffon’s needle images at different lines’ densities and calculating the variance for different angles, shows a clear marker for symmetry breaking and loss of randomness in these images throughout the entire lines’ density range (SI, Fig S11).

Since the Radon transform can be used to estimate line lengths and thence address Bertrand’s problem, we created 100 images for each method (*C, R, A*) and for each chord density (2,100 images in total, see Methods), calculated its Radon transform, and summed over angles to get the marginal Radon transform for each line length. From these probability distributions it is straightforward to calculate the probability for a random chord to be longer than the triangle’s side (Fig 2e). We see that Radon transform estimates of Bertrand’s probability preserve the ranking of the different methods *P*_*R*_ *> P*_*C*_ *> P*_*A*_, similar to participants’ estimates and with a similar range to that of the behavioral results, though with opposing monotonicity (Fig 3c).

Our study has explored the use of geometric probability to assess the visual estimation of randomness and probability, by exploiting its link to spatial probabilistic settings and invariance properties. Our results suggest that people perceive images that hold more invariances (out of the subset of rotational, translational, and scale invariances) as more random even at low chord densities where geometrical information is scarce.

How do people then quantify randomness and probability from images? At the neuronal level, for these computations to be plausible, we need the ability to measure orientations (for Buffon’s needle) and lengths (for Bertrand’s paradox). There is evidence for how the visual cortex uses basic neural mechanisms to carry out computations of orientations and lengths. Simple cells neurons in the visual cortex have long been known to be able to discriminate orientational order [31, 32], with recent evidence for how this is embedded in neuronal geometry and firing rates [33]. Similarly, end-stopped simple and complex cells in the visual cortex can signal length of lines at different orientations [34–37]. Thus, the response of these neurons provide the plausible first layer for the computations linking probability, randomness, and symmetry.

At the cortical level, the visual system needs to integrate the signals from simple and complex cells to infer orientation, translation and scale invariances. The Radon transform, a fundamental object in integral geometry that is invariant under translation, rotation and scale is a natural candidate for assessing symmetry, randomness and probability in images of planar lines. Given the strong correspondence between our Radon transform analysis and participants’ behavioral results, combined with the fact that length processing resides mainly in the intraparietal sulci (IPS) [38–42], and the possibility of a Radon transform implementation in visual neural networks [43–46], it is tempting to search for a neuronal circuit in the human brain that integrates line directions, lengths and their propensity in visual stimuli.

In an uncertain world, estimating randomness and probability provides a strong selective advantage, particularly in visual perception. Visual processing requires geometrical notions and takes the form of local orientational correlations which has been well-established. Our study suggests that we are equally capable of deducing non-local properties of images and assess randomness and probability. Integral geometry provides a natural language for the consideration of global geometrical questions in a statistical setting. This study emphasizes these notions by demonstrating how perception of randomness depends at both the geometrical and physical attributes of the problem. It appears that people’s global inference in the visual domain of geometric probability is quite robust, consistent with the deep mathematical link between probability, randomness, and invariance.

## Acknowledgments

We thank the Harvard Inter-faculty Mind, Brain and Behavior initiative for partial financial support, and Avraham E. Mayo for fruitful discussions.

*

